# Early Patellofemoral Cartilage and Bone Degeneration in a Rat Model of Noninvasive Anterior Cruciate Ligament Rupture

**DOI:** 10.1101/2021.04.11.439337

**Authors:** Samantha E. Hartner, Michael D. Newton, Mackenzie M. Fleischer, Kevin C. Baker, Tristan Maerz

## Abstract

**Background:** Anterior cruciate ligament rupture (ACLR) is a well-known risk factor for the development of post-traumatic osteoarthritis (PTOA). While clinical and pre-clinical studies have characterized the onset and progression of PTOA in the tibiofemoral joint compartment, very little is known about degenerative changes in the patellofemoral compartment after ACL injury.

**Hypothesis/Purpose:** To evaluate the extent to which ACL rupture induces acute patellofemoral joint degeneration by quantifying articular cartilage morphology and remodeling of subchondral and trabecular bone microarchitecture in the patellofemoral compartment.

**Study Design:** Descriptive laboratory study.

**Methods:** Adult female Lewis rats were randomized to undergo either a non-surgical ACL rupture or a Sham procedure (n = 6 per group). *Ex vivo* contrast-enhanced micro-computed tomography (µCT) and histological evaluation of the patellofemoral compartment were performed at 2-weeks post-injury, representing a timepoint of documented early PTOA in the tibiofemoral compartment in this model.

**Results:** ACL rupture causes osteophyte formation in the patella and mild degeneration in the superficial zone of articular cartilage (AC), including surface fibrillation, fissures, increased cellularity, and abnormal chondrocyte clustering at two weeks post-injury. Contrast-enhanced µCT analysis demonstrates significant increases in AC thickness of patellar and trochlear cartilage. Loss of subchondral bone thickness, bone volume fraction, and tissue mineral density, as well as changes to trabecular microarchitecture in both the patella and trochlea, were indicative of catabolic bone remodeling.

**Conclusion:** These results demonstrate that the patellofemoral joint develops mild but evident degenerative changes in the acute time period following ACL rupture, extending the utility of this rat model to the study of degenerative patellofemoral changes following joint trauma.

**Clinical Relevance:** ACL rupture causes mild degeneration and swelling of articular cartilage, coupled with catabolic bone remodeling in the patellofemoral compartment. Characterizing the pathophysiology of patellofemoral PTOA in its early stages may provide a better understanding of disease progression and provide opportunities for preventative therapeutic intervention.

## INTRODUCTION

Anterior cruciate ligament (ACL) rupture is one of the most common soft-tissue musculoskeletal injuries and a well-known risk factor for post-traumatic osteoarthritis (PTOA)^4,24^. The progression of tibiofemoral PTOA following ACL rupture has been well-characterized in both clinical and pre-clinical literature^6,22,25,41^. However, comparatively few studies have characterized patellofemoral compartment OA (PFOA) following ACL rupture, and relatively little is known regarding incidence and overall phenotypic manifestation of joint degeneration in this compartment. Recent work suggests that PFOA may be equally or even more prevalent than tibiofemoral OA: Culvenor *et al* demonstrate that within 5-10 years post-ACL rupture, 47% of patients exhibit PFOA progression – including osteophytes, cartilage lesions, and joint space narrowing – compared to a 31% incidence of tibiofemoral OA^9^. Furthermore, recent MRI-based assessment of PFOA indicated that 40% of patients exhibit detectable patellofemoral cartilage loss within 5 years of ACL injury, with cartilage thickness in the trochlea decreasing by as much as 1% annually in the first 2 years^8^.

Despite clinical evidence of a high incidence of PFOA development after ACL injury, the acute manifestations of PFOA and the associated microstructural, cellular, and molecular changes following ACL injury remain poorly understood. Clinical evidence regarding the nature of early patellofemoral tissue changes is lacking, as most clinical studies of PFOA track joint degeneration once disease has already developed. Natural history studies assessing PFOA incidence and development commonly employ radiography^9,20,32,34^, which can diagnose joint space narrowing and osteophyte formation but is inherently incapable of assessing the extent and location of articular cartilage and bone degeneration, especially in early disease. While MRI is the gold-standard for assessment of cartilage thickness and bone marrow lesions^7,8,21^, it lacks imaging resolution to measure injury-induced changes to bone microarchitecture. Thus, the collective understanding of acute PFOA following ACL injury is currently limited.

Small animal models of ACL injury have become invaluable tools for the preclinical study of PTOA. In a previous rat study of surgical ACL transection, decreased subchondral bone (SCB) perfusion and associated cartilage and bone lesions were observed in the patellofemoral compartment at intermediate and chronic time points following injury^40^, demonstrating chronic manifestations of ACL injury in the rodent patellofemoral compartment. However, this study did not analyze acute time points to understand early disease, and surgical models of ACL injury are inherently limited in their assessment of acute injury due to the confounding influence of surgical trauma and lack of mechanical loading. Non-invasive models of isolated ACL rupture, whereby joint injury is induced mechanically, provide a highly controlled and clinically-representative means of inducing joint injury, and the utilization of these models to study tibiofemoral PTOA has increased in recent years^3,6,23,26,38^. However, to date, the extent of patellofemoral remodeling and degeneration in these non-invasive injury models remains largely unexplored, and it is unclear to what extent these models may be employed to study PFOA pathophysiology and potential treatments. To this end, the aim of this study was to employ both quantitative imaging and histological evaluation to characterize acute changes to bone and articular cartilage within the patellofemoral joint in a rat model of non-invasive ACL rupture.

## MATERIALS AND METHODS

### Rats and Non-Invasive ACL Rupture

Under an IACUC-approved protocol, 12 female Lewis rats (aged 14 weeks, weight ∼200-220g, Charles River Laboratories, Wilmington, MA) were randomized to either non-invasive ACL rupture (ACLR) or Sham groups (*n* = 6 per group). The sample size was determined based on published μCT data from the same model^25^: given the previously-observed effect size in trabecular bone volume fraction between control rats and ACLR rats (a morphometric parameter most responsive to injury and an outcome we deemed most applicable to the present study) in order to detect a 5% difference between groups, (effect size = 2.2, α = 0.05, power = 0.9), six rats would be required. To induce ACLR or Sham injury, rats were placed prone on a materials testing system (Insight 5, MTS, Eden Prairie, MN) with the right knee positioned in 100° of flexion and the hindpaw held in 30° of dorsiflexion in a custom fixture. In the ACLR group, ACL rupture was induced by applying a preload, ten preconditioning cycles, and a subsequent rapid compressive displacement to the knee joint, as described previously^26^. The Sham group was subjected to preload and preconditioning cycles only. Anesthesia was induced by intraperitoneal ketamine and xylazine, and maintained under 1%-2% inhaled isoflurane. All rats were administered 5 mg/kg subcutaneous carprofen immediately prior to Sham/ACLR injury, and buprenorphine (1.2 mg/kg) was administered for analgesia at 12h and 24h post-procedure. All procedures were completed in the morning, and rats individually housed and allowed *ad libitum* cage activity and access to standard rodent chow under a 12-hour light/dark cycle for 2 weeks post-procedure until euthanasia via CO_2_ asphyxiation. To perform contrast-enhanced imaging, femora and patellae were completely dissected to expose the articular surface, fixed in 10% neutral buffered formalin for 48 hours, and transferred to 70% ethanol for long-term storage.

### Ex Vivo Micro-Computed Tomography

To facilitate contrast-enhanced µCT of articular cartilage, femora and patellae were rehydrated in PBS overnight and incubated in 20% v/v ioxaglate (Hexabrix 320, Guerbet LLC, Princeton, NJ) for 2 hours prior to µCT imaging as previously described^36^. In preliminary experiments, a 2-hour contrast agent incubation was found to be sufficient to achieve equilibrium in articular cartilage (data not shown). Both patellae and femora were securely positioned in custom sample holders for consistent imaging positioning and orientation, and humidifying silica beads were used to maintain a 70% humid environment during µCT acquisition. Femora were scanned on a µCT-40 (Scanco Medical AG, Brüttisellen, Switzerland) at 55 kVp, 145 µA, 300 ms integration time, with an isotropic voxel size of 8 µm. Patellae were scanned on a VivaCT-80 (Scanco Medical AG) at 55 kVp, 145 µA, 800 ms integration time, with a 12 µm isotropic voxel size. The use of two imaging systems was necessary given study timeline restrictions, and validation data demonstrates equivalent morphometry results between these two imaging resolutions (data not shown). Furthermore, no comparisons are made between the two imaging protocols. Following scanning, specimens were washed in PBS for 30 minutes and graded back to 70% ethanol.

### Imaging Analysis

Femoral and patellar articular cartilage, subchondral bone, and trabecular bone volumes of interest were isolated using an automated, atlas registration-based segmentation algorithm in MATLAB (Mathworks, Natick, MA), as previously described^33^. All segmentations were reviewed for accuracy by an experienced investigator. Femoral AC and bone volumes were further segmented to isolate the trochlear compartment, defined as all bone and AC proximal to the femoral notch. AC was quantitatively analyzed via mean thickness (AC.Th) using the direct-distance transform (i.e “sphere-fitting”) method in boneJ^11^. Trabecular bone was assessed via bone volume fraction (BV/TV), tissue mineral density (TMD), trabecular thickness (Tb.Th), trabecular spacing (Tb.Sp), and trabecular number (Tb.N) according to standard methods^2^. Subchondral bone was assessed via subchondral bone thickness (SCB.Th), BV/TV, and TMD. A standardized hydroxyapatite phantom was used to calibrate imaging data to mineral density.

### Histology

Patellae, including the attached quadriceps and patellar tendons, were decalcified, embedded in paraffin and bisected sagittally, and a single 5 µm sagittal section (including both halves) was taken. Femora underwent additional histological assessments of the femoral condyles as part of a separate study on the tibiofemoral compartment which required the use of non-decalcified methods^28^. As such, femora were embedded in methyl methacrylate and bisected into medial and lateral halves. Two 5 µm sections were taken, spaced 200 µm apart. For this study, trochlear analysis was carried out on lateral compartment sections, which fully encompassed the trochlear articular cartilage. All sections of both patellae and femora were stained with Safranin-O/Fast Green and imaged at 20x magnification using a slide imaging system (Aperio, Leica 122 Biosystems, Buffalo Grove, IL). Two blinded raters scored each section using the Osteoarthritis Research Society International (OARSI) Modified Mankin score^16^, and graders conferred on disparate scores (differences of greater than one point) until consensus was reached. Histologic scores were averaged across all sections for each limb, and an aggregate group average was calculated.

### Statistical Analyses

Statistical analysis was performed using SPSS (IBM, Armonk, NY). The normality and equal variance assumptions were confirmed for continuous data. µCT data were assessed via mixed two-way analysis of variance (ANOVA) with one within-subject factor (Injured vs Contralateral) and one between-subject factor (ACLR vs Sham). Histologic grades were assessed via the Kruskal Wallis test, with *post hoc* comparisons via Mann-Whitney U tests with a Bonferroni correction for multiple comparisons. Adjusted P values below 0.05 were considered significant.

## RESULTS

All animals maintained good health status for the 2-week period following the ACLR and Sham procedures and thus all 6 rats from both experimental groups were included in the analysis.

### Articular Cartilage Exhibits Acute Swelling Following ACLR

AC was significantly thicker in the ACLR Injured limb compared to the Contralateral limb in the patella and trochlea (Patella – 12.3% increase, *P* = 0.003; Trochlea – 8.7% increase, *P* = 0.005), and compared to the Sham Uninjured limb in the patella (18.8% increase, *P* = 0.001) (**Figure 1**). Interestingly, patellar AC of the Sham Uninjured limb was significantly thinner compared to the Sham Contralateral (8.1% decrease, *P* = 0.026), indicating a subtle effect of sham loading. In the patella of the ACLR Injured limb, AC thickening occurred over the entire superior cartilage surface, encompassing both the medial and lateral regions, while in the Sham Uninjured limb, cartilage thinning was most pronounced on the superior and lateral surfaces (**Figure 1A**). In the trochlea of the ACLR Injured limb, AC thickening occurred primarily in the proximal aspect but with reduced magnitude compared to the patella (Trochlea: 8.7% increase; Patella: 12.3% increase). Taken together, these findings demonstrate that ACLR induces acute swelling/thickening of both patellar and trochlear articular cartilage, with a greater magnitude of change observed in the patella.

**Figure 1.**
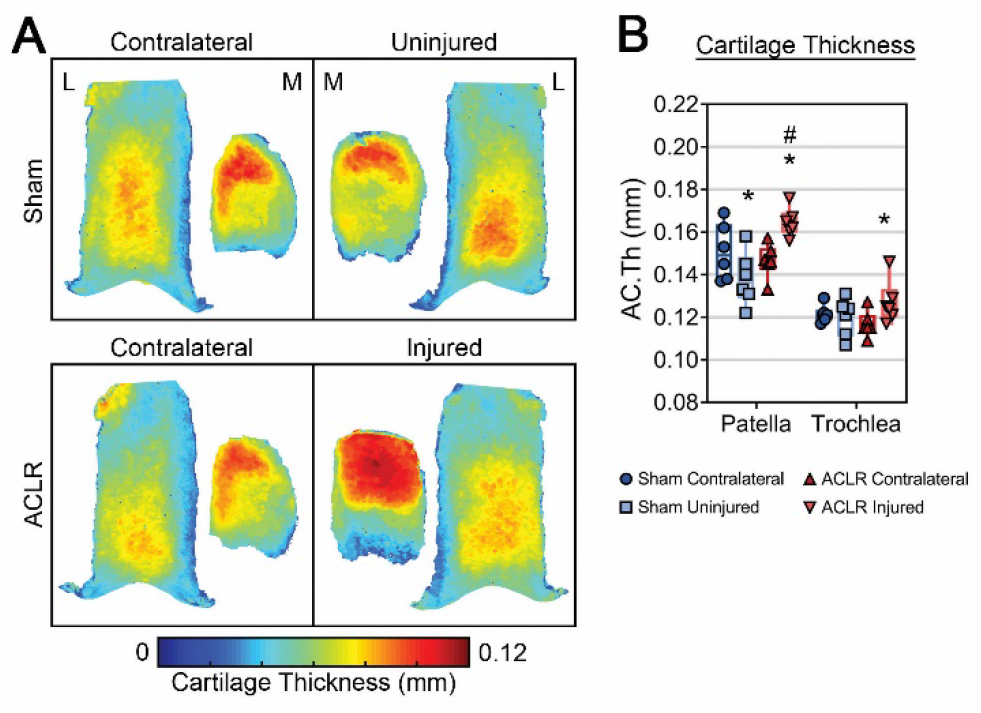
Three-dimensional articular cartilage thickness maps (A) and numerical thickness results (B) derived from contrast-enhanced µCT demonstrate thickening of both patellar and trochlear cartilage following ACLR. The top of the figure represents the superior/proximal aspect of the patella and trochlea. n = 6 for all groups. * represents significant difference compared to Contralateral limb (^*^P < 0.05, ^**^P < 0.001); # represents significant difference compared to Sham Uninjured limb (^#^P < 0.05, ^##^P < 0.001); mixed two-way ANOVA.

### Injury-Induced Loss of Trabecular and Subchondral Bone is Observed in the Patella and Trochlea

ACLR induced catabolic trabecular and subchondral bone remodeling of both the patella and the trochlea. In the patella, trabecular BV/TV, TMD, Tb.Th, and Tb.N were significantly decreased and Tb.Sp was significantly increased in the ACLR Injured limb compared to the Contralateral limb (BV/TV: 12.9% decrease, *P* < 0.001, TMD: 2.3% decrease, *P* = 0.001, Tb.Th: 6.8% decrease, *P* < 0.001, Tb.N: 6.5% decrease, *P* < 0.001, Tb.Sp: 10.1% increase, *P* < 0.001) and compared to the Sham Uninjured limb (BV/TV: 12.7% decrease, *P* < 0.001, TMD: 2.0% decrease, *P* = 0.046, Tb.Th: 6.4% decrease, *P* = 0.022, Tb.N: 6.8% decrease, *P* = 0.001, Tb.Sp: 10.8% increase, *P* < 0.001) (**Figure 2**). In the trochlea, epiphyseal Tb.Th was significantly decreased in the ACLR Injured limb compared to both the Contralateral (4.3% decrease, *P* = 0.008) and Sham Uninjured limbs (6.2% decrease, *P* = 0.002). Epiphyseal BV/TV was significantly lower in the ACLR Injured limb compared to the Sham Uninjured limb (7.3% decrease, *P* = 0.023). In the subchondral compartment, subchondral BV/TV and thickness were significantly decreased in the ACLR Injured limb compared to the Contralateral limb (Patella - BV/TV: 3.6% decrease, *P* = 0.001, SCB.Th: 12.0% decrease, *P* < 0.001; Trochlea - BV/TV: 2.5% decrease, *P* < 0.001, SCB.Th: 11.5% decrease, *P* < 0.001) and compared to the Sham Uninjured limb (Patella - BV/TV: 4.2% decrease, *P* < 0.001, SCB.Th: 15.0% decrease, *P* = 0.002; Trochlea - BV/TV: 2.9% decrease, *P* = 0.002, SCB.Th: 15.7% decrease, *P* = 0.001) (**Figure 2**). Although no formal region-dependent analysis was performed, loss of SCB.Th appeared to occur globally across the patella and trochlea in the ACLR limb, while in the Sham limb, increased SCB.Th appears more pronounced on the inferior patella and medial trochlea. Patellar SCB TMD was significantly decreased in the ACLR Injured limb compared to the Contralateral limb (3.0% decrease, *P* < 0.001). SCB.Th of the Sham Uninjured limb was significantly higher compared to the Sham Contralateral limb (Patella: 5.1% increase, *P* = 0.014, Trochlea: 5.6% increase, *P* = 0.010). Taken together, these results demonstrate the acute loss of both trabecular and subchondral bone in the patellofemoral compartment following ACLR.

**Figure 2.**
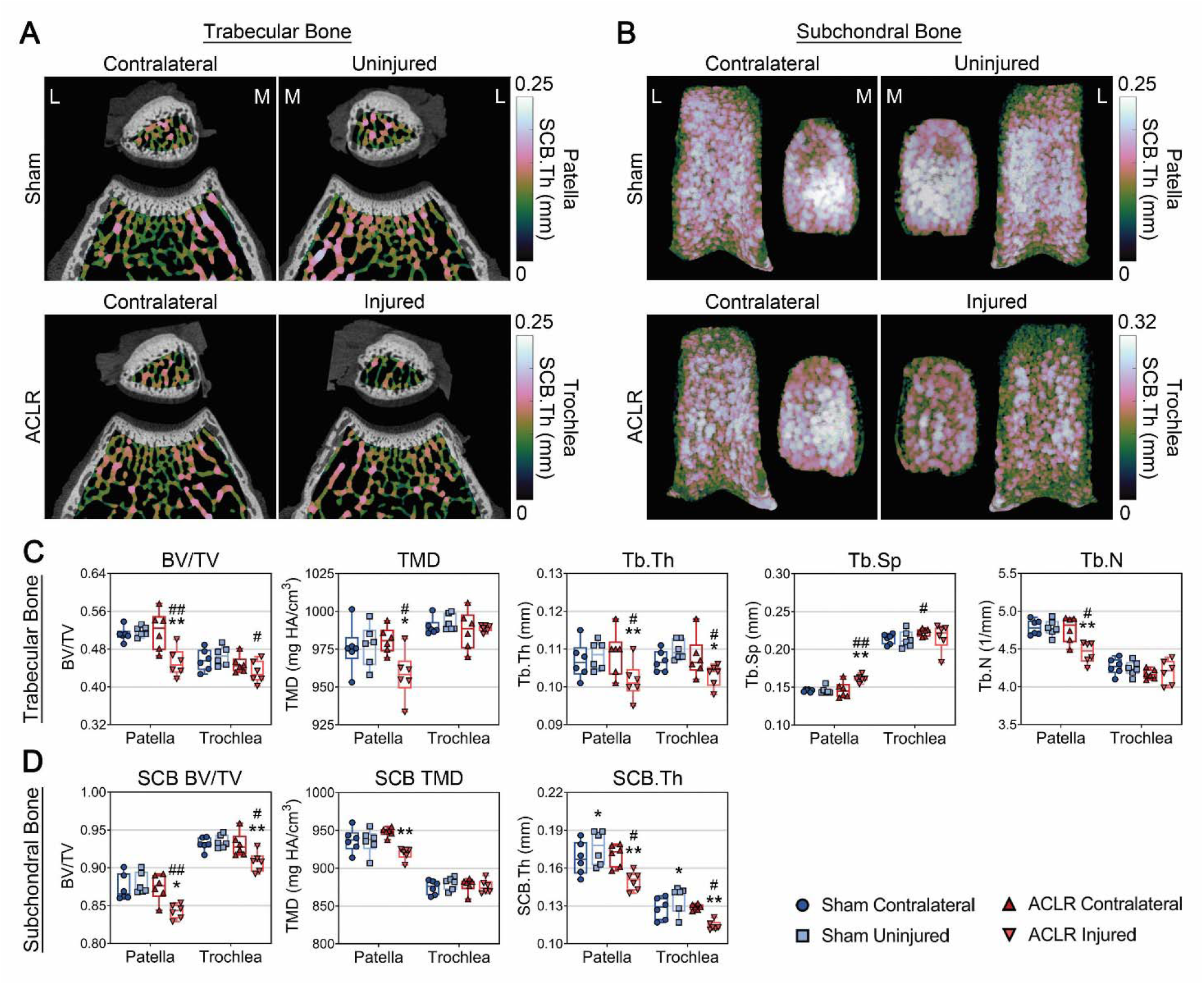
Trabecular bone thickness maps of the patella and trochlea overlaid onto representative axial µCT images show increased Tb.Sp and reduced Tb.Th and Tb.N in the ACLR Injured limb (A). 3D thickness maps show reduced subchondral bone thickness in the patella and trochlea (B). Morphometric results demonstrate catabolic remodeling of both trabecular (C) and subchondral bone (D) following ACL rupture. The overall magnitude of catabolic remodeling was greater in the patella compared to the trochlea. The top of the figure in B represents the superior/proximal aspect of the patella and trochlea. n = 6 for all groups. * represents significant difference compared to Contralateral limb (^*^P < 0.05, ^**^P < 0.001); ^#^ represents significant difference compared to Sham Uninjured limb (^#^P < 0.05, ^##^P < 0.001); mixed two-way ANOVA.

### Histology Demonstrates Superficial Articular Cartilage Damage, Osteophyte Formation, and Abnormal Chondrocyte Morphology Following ACL Rupture

Patellar and trochlear cartilage of both Sham limbs appeared normal, with a smooth articular surface, normal chondrocyte morphology, and homogenous Saf-O staining distributions. While our contrast-enhanced μCT results demonstrate mild thinning of Sham Uninjured cartilage in the patella, we did not observe any notable degenerative changes in Sham limbs on histology. ACLR Injured patellae demonstrate mild but evident degenerative histological changes, including surface fibrillation, reduction of intracellular and extracellular PG staining, increased chondrocyte clustering, and osteophyte formation (**Figure 3A-B**). We also observed an increased incidence of hypertrophic chondrocytes in deep cartilage, most notably at the superior patellar aspect where articular cartilage thickening was observed. In superficial patellar cartilage, a greater incidence of abnormal chondrocyte clustering was noted in ACLR Injured limbs (**Figure 3A, red arrowheads**). Superficial fissures and abnormal chondrocyte clustering were noted to be more prevalent in the inferior region of patellar cartilage. Osteophytes appeared immature, with primarily cartilaginous composition and focal regions of mineralized tissue, indicative of active osteophyte formation. This osteophyte formation was exclusively observed at the superior aspect of the patella (**Figure 3B**). Correspondingly, ACLR Injured patellae had a significantly higher OARSI score compared to both ACLR Contralateral (*P* = 0.015) and Sham Uninjured limbs (*P* = 0.015), as well as several significantly higher sub-scores compared to both Contralateral (AC Structure: *P* = 0.018, PG Staining: *P* = 0.014, Cellularity: *P* = 0.008, Osteophyte: *P* = 0.004) and Sham Uninjured limbs (AC Structure: *P* = 0.014, PG Staining: *P* = 0.014, Cellularity: *P* = 0.007, Osteophyte: *P* = 0.004) (**Figure 3C**).

**Figure 3.**
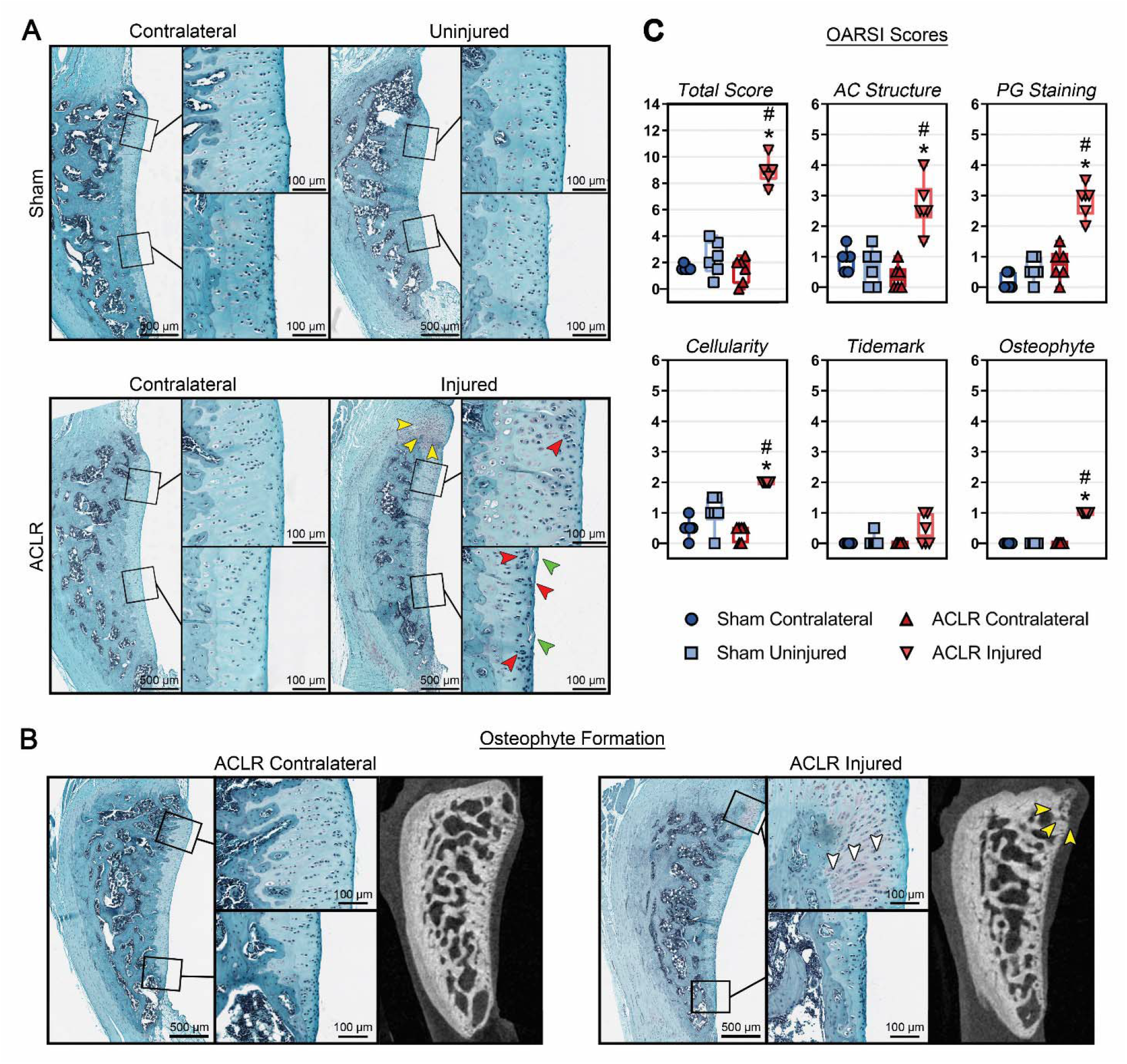
Representative sagittal histology of patellae demonstrates that ACLR Injured patellae exhibit superficial fissures (green arrowheads), osteophyte formation (yellow arrowheads), and abnormal chondrocyte morphology and clustering (red arrowheads) (A). Osteophyte formation at the superior aspect of ACLR Injured patellae was evident on both histology and µCT. Proliferation of hypertrophic and pre-hypertrophic chondrocytes indicative of endochondral ossification is evident in superior aspect of the ACLR Injured patella (white arrowheads) (B). ACLR Injured patellae exhibited significantly higher OARSI Modified Mankin scores and sub-scores compared to both ACLR Contralateral and Sham Uninjured patellae (Maximum OARSI scores: Total Score – 21; AC Structure – 8; PG Staining – 6; Cellularity – 3; Tidemark – 1; Osteophyte – 3) (C). The top of the figure represents the superior/proximal aspect of the patella. n = 6 for all groups. * represents significant difference compared to Contralateral limb (P < 0.05); ^#^ represents significant difference compared to Sham Uninjured limb (P < 0.05); Kruskal-Wallis ANOVA.

ACLR Injured trochleae had largely intact cartilage surfaces, with only minor noted superficial fibrillation (**Figure 4A, green arrowhead**). Notably, ACLR trochleae exhibited focal regions of intense Saf-O staining and pockets of abnormal clustering/cloning in superficial-layer cells. ACLR Injured trochleae had a significantly higher OARSI score compared to Sham Uninjured limbs (*P* = 0.024). Multiple OARSI sub-scores were significantly higher in ACLR trochleae compared to Sham Uninjured limbs (AC Structure: *P* = 0.038, PG Staining: *P* = 0.022, Cellularity: *P* = 0.023) and the tidemark sub-score was significantly higher compared to ACLR Contralateral limbs (*P* = 0.035) (**Figure 4B**). Taken together, our histologic results demonstrate that at this acute post-injury timepoint, ACL rupture induces mild articular cartilage degeneration of the patella and trochlea, largely contained to the superficial zone, and osteophyte formation at the superior aspect of the patella.

**Figure 4.**
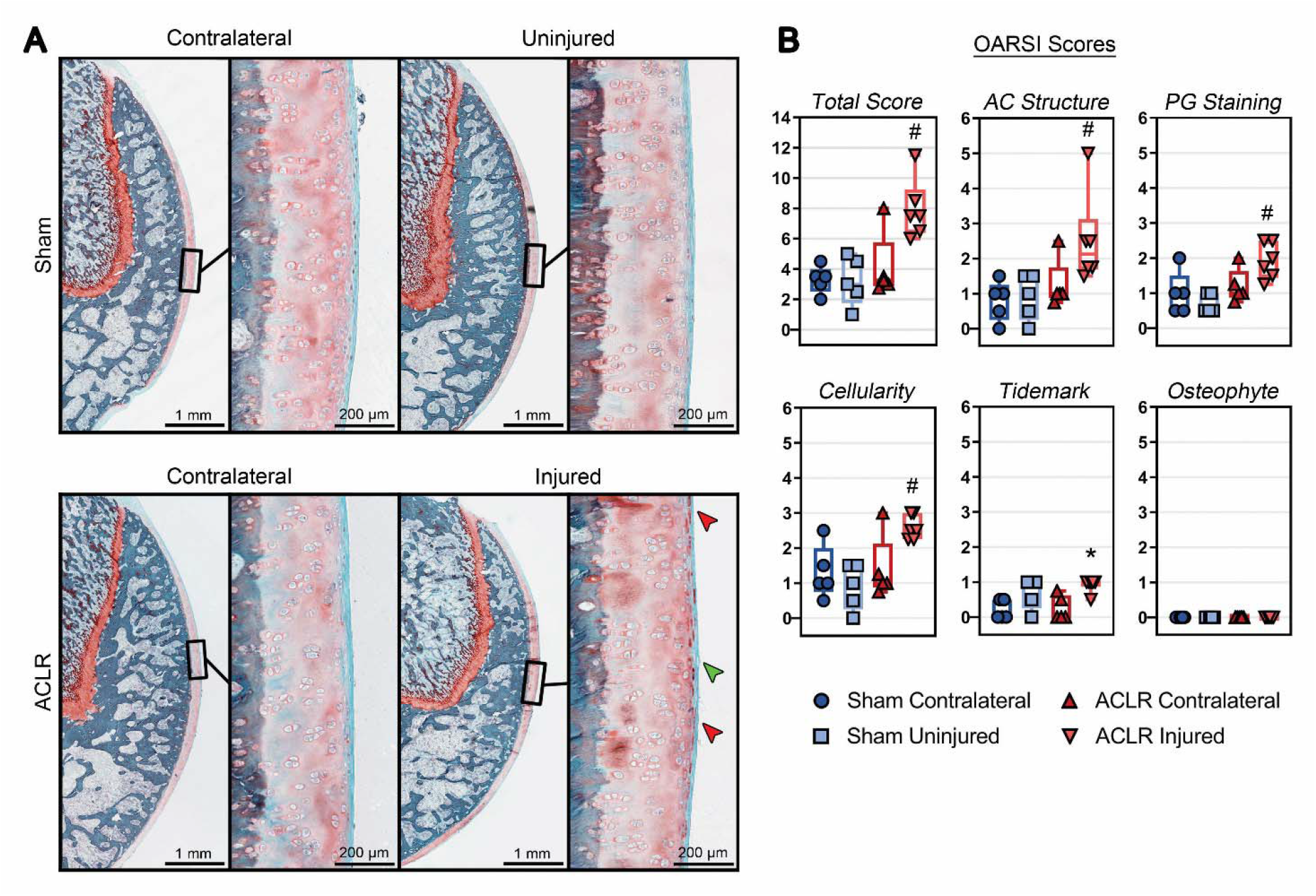
Representative sagittal trochlea histology shows that ACLR Injured patellae exhibit mild superficial fissures (green arrowhead), and abnormal chondrocyte morphology, clustering, and focal regions of intense PG staining (red arrowheads) (A). ACLR Injured trochleae exhibit significantly higher OARSI Modified Mankin scores and sub-scores compared to Sham Uninjured trochleae (Maximum OARSI scores: Total Score – 21; AC Structure – 8; PG Staining – 6; Cellularity – 3; Tidemark – 1; Osteophyte – 3) (B). The top of the figure represents the superior/proximal aspect of the trochlea. n = 6 for all groups. * represents significant difference compared to Contralateral limb (P < 0.05); ^#^ represents significant difference compared to Sham Uninjured limb (P < 0.05); Kruskal-Wallis ANOVA.

## DISCUSSION

Significant articular cartilage and subchondral bone remodeling in the weight-bearing regions of the tibiofemoral compartment are the hallmarks of PTOA in the knee joint. However, comparatively little is known about the development and progression of PTOA in the patellofemoral compartment following ACL injury, or how PFOA contributes to overall disease progression in the knee joint. In this study we aimed to characterize the onset of patellofemoral cartilage and bone degeneration in a rat model of non-invasive ACL rupture, in which changes to the patellofemoral compartment have not been characterized to date. Our results demonstrate catabolic bone remodeling in both subchondral and trabecular bone, in addition to articular cartilage swelling, abnormal changes to chondrocyte morphology and clustering, patellar osteophyte formation, and surface degeneration of articular cartilage. Collectively, these findings support that ACL rupture induces early patellofemoral compartment degeneration in this rat model.

At 2 weeks following ACLR, quantitative µCT revealed significant thickening/swelling of the articular cartilage in both the patella and trochlea while histological assessment showed signs of superficial damage including surface fibrillation and fissuring. We observed an increased incidence of hypertrophic chondrocytes in the deep AC layer of injured joints, which qualitatively co-localized with regions of AC thickening observed in µCT. Taken together, these manifestations are indicative of early cartilage disease. Studies in both humans and animal models have shown that PTOA-associated AC degeneration initiates in the superficial zone with the development of surface irregularities, fissures, and disruption of the superficial collagen matrix^17,18,39^. Regarding cartilage thickness changes in response to injury, while some clinical studies of PTOA with early MRI assessment report compartment-specific regions of AC thickening^14,27,29^, other studies report short-term cartilage thickness loss that often persists into chronic disease^8,13,37^. Previous studies of ACL rupture in rats have reported AC thickening in the tibial plateau, femoral condyles, and trochlea persisting for up to 10 weeks post-injury^29^, but to our knowledge this is the first study to report acute AC thickening in the patella using high-resolution µCT. In contrast to our findings, a recent study employing ACL transection in rats reported that both patellar and trochlear AC were histologically intact at 16 weeks post-injury, with surface degeneration and hypertrophic chondrocytes presenting by 32 weeks^40^. While this may reflect inherent differences between surgical vs non-invasive ACL injury, our findings of early/mild cartilage damage underscore the value of high-resolution µCT in measuring the initial cartilage response to traumatic injury, which may be undetected and cannot be comprehensively quantified using histology or lower-resolution imaging modalities.

Acute loss of bone mineral density and bone volume following ACL rupture have been well-described in clinical literature^22,41^. Both clinical and pre-clinical studies have found that in the short term following ACL injury, subchondral and trabecular bone undergo catabolic degeneration, followed by a transition toward anabolic remodeling. In the subchondral bone, this remodeling persists in the long term, leading to subchondral bone sclerosis^1,6,21^. While animal models have accurately characterized this remodeling using high-resolution imaging, the exact nature of injury-induced catabolic bone remodeling in humans remains undescribed given that, to date, there have been very few clinical studies employing high-resolution imaging capable of accurately measuring bone microarchitecture. Recently, however, Kroker *et al* used HR-pQCT to demonstrate that femoral subchondral bone thickness and BMD decreases for 7-8 months post-ACL injury before entering a plateau or recovery phase^21^. The subchondral and trabecular bone results of the current study share key similarities with this clinical work, affirming that at 2-weeks following ACLR, distinct bone remodeling-related indicators of early PFOA are readily observable in this rat model.

A notable finding in our dataset is that several injury-induced changes occurred with greater magnitude in the patella compared to changes in the trochlea. While a few prior clinical studies have reported more severe cartilage loss in the patella compared to the femoral condyles^37^ and trochlea^35^, at least one other study has found that greater cartilage degeneration occurs in the trochlea in the 2-5 year post-injury time period^8^. Thus, it remains unclear to what extent this finding is specific to this animal model or specific to a stage of PTOA severity, and it is beyond the scope of the current study to investigate the cause of this difference. However, previous clinical research has demonstrated that compared to the other articular surfaces of the knee, patellar AC uniquely exhibits a differential compressive strain response to variations in physical activity and intensity level, suggesting that the patella may be more sensitive to mechanical stimuli and instability than the trochlea^12,35^. This is consistent with cadaveric findings that patellar AC has higher fluid permeability and lower proteoglycan content than femoral AC, potentially making the patella more susceptible to mechanically-induced degeneration^15^. It is also possible that the mechanism of injury used in our model places a load on the patella during compression and subluxation of the tibia. Given the location of the patella relative to the trochlea in 100° of flexion, this compression could be inducing some mild degeneration in the patella while having a lesser effect on the trochlea. Future studies may assess biomechanical and molecular mechanisms underpinning differential disease severity in the two surfaces of the patellofemoral compartment.

Some limitations should be taken into consideration when interpreting the results of this study. Although the Sham loading procedure did not cause any macroscopic injury or histologically-observable degeneration, it did have measurable effects on the AC and subchondral bone of the Sham limb compared to its contralateral. Sham effects are frequently reported in other models of rodent ACL injury, including patellar tendon ossification, increased AC water content, synovitis, and mild loss of proteoglycan at the AC surface resulting from a sham surgical procedure^5,19,30,31^. The non-invasive Sham procedure described here may have benefits over a surgical procedure involving skin and muscle incision and arthrotomy, which could introduce multiple biological confounding effects. Finally, due to the differences in histological processing and µCT scanning protocols used for the patella and the femur, we did not make any direct comparisons between the patella and trochlea.

In this rat model of non-invasive ACL rupture, we observed acute degeneration of the patellofemoral compartment, characterized by cartilage thickening and mild articular surface damage alongside catabolic trabecular and subchondral bone remodeling. Injury-induced changes in articular cartilage and trabecular bone were of greater severity in the patella than changes in the trochlea. These results demonstrate the acute onset of mild patellofemoral joint degeneration following ACL rupture and extend the utility of this rat joint injury model to the study of patellofemoral PTOA.

## Supporting information

IACUC Approval

ARRIVE Checklist

## REFERENCES

1. Anderson DD, Chubinskaya S, Guilak F, et al. Post-traumatic osteoarthritis: improved understanding and opportunities for early intervention. J Orthop Res. 2011;29(6):802–809.

2. Bouxsein ML, Boyd SK, Christiansen BA, et al. Guidelines for assessment of bone microstructure in rodents using micro-computed tomography. J Bone Miner Res. 2010;25(7):1468–1486.

3. Brown SB, Hornyak JA, Jungels RR, et al. Characterization of Post□Traumatic Osteoarthritis in Rats Following Anterior Cruciate Ligament Rupture by Non□Invasive Knee Injury (NIKI). J Orthop Res. 2019.

4. Carbone A, Rodeo S. Review of current understanding of post-traumatic osteoarthritis resulting from sports injuries. J Orthop Res. 2017;35(3):397–405.

5. Chou M-C, Tsai P-H, Huang G-S, et al. Correlation between the MR T2 value at 4.7 T and relative water content in articular cartilage in experimental osteoarthritis induced by ACL transection. Osteoarthritis Cartilage. 2009;17(4):441–447.

6. Christiansen BA, Anderson MJ, Lee CA, et al. Musculoskeletal changes following non-invasive knee injury using a novel mouse model of post-traumatic osteoarthritis. Osteoarthritis Cartilage. 2012;20(7):773–782.

7. Culvenor AG, Collins NJ, Guermazi A, et al. Early Patellofemoral Osteoarthritis Features One Year After Anterior Cruciate Ligament Reconstruction: Symptoms and Quality of Life at Three Years. Arthritis Care Res (Hoboken). 2016;68(6):784–792.

8. Culvenor AG, Eckstein F, Wirth W, Lohmander LS, Frobell R. Loss of patellofemoral cartilage thickness over 5 years following ACL injury depends on the initial treatment strategy: results from the KANON trial. Br J Sports Med. 2019;53(18):1168–1173.

9. Culvenor AG, Lai CC, Gabbe BJ, et al. Patellofemoral osteoarthritis is prevalent and associated with worse symptoms and function after hamstring tendon autograft ACL reconstruction. Br J Sports Med. 2014;48(6):435–439.

10. Culvenor AG, Segal NA, Guermazi A, et al. Sex□Specific influence of quadriceps weakness on worsening Patellofemoral and Tibiofemoral cartilage damage: a prospective cohort study. Arthritis Care Res. 2019;71(10):1360–1365.

11. Doube M, Kłosowski MM, Arganda-Carreras I, et al. BoneJ: free and extensible bone image analysis in ImageJ. Bone. 2010;47(6):1076–1079.

12. Eckstein F, Lemberger B, Gratzke C, et al. In vivo cartilage deformation after different types of activity and its dependence on physical training status. Ann Rheum Dis. 2005;64(2):291–295.

13. Eckstein F, Maschek S, Wirth W, et al. One year change of knee cartilage morphology in the first release of participants from the Osteoarthritis Initiative progression subcohort: association with sex, body mass index, symptoms and radiographic osteoarthritis status. Ann Rheum Dis. 2009;68(5):674–679.

14. Frobell RB. Change in cartilage thickness, posttraumatic bone marrow lesions, and joint fluid volumes after acute ACL disruption: a two-year prospective MRI study of sixty-one subjects. J Bone Joint Surg Am. 2011;93(12):1096–1103.

15. Froimson MI, Ratcliffe A, Gardner TR, Mow VC. Differences in patellofemoral joint cartilage material properties and their significance to the etiology of cartilage surface fibrillation. Osteoarthritis Cartilage. 1997;5(6):377–386.

16. Gerwin N, Bendele A, Glasson S, Carlson CS. The OARSI histopathology initiative– recommendations for histological assessments of osteoarthritis in the rat. Osteoarthritis Cartilage. 2010;18:S24–S34.

17. Guilak F, Ratcliffe A, Lane N, Rosenwasser MP, Mow VC. Mechanical and biochemical changes in the superficial zone of articular cartilage in canine experimental osteoarthritis. J Orthop Res. 1994;12(4):474–484.

18. Hollander AP, Pidoux I, Reiner A, et al. Damage to type II collagen in aging and osteoarthritis starts at the articular surface, originates around chondrocytes, and extends into the cartilage with progressive degeneration. J Clin Invest. 1995;96(6):2859–2869.

19. Jackson MT, Moradi B, Zaki S, et al. Depletion of protease□activated receptor 2 but not protease□activated receptor 1 may confer protection against osteoarthritis in mice through extracartilaginous mechanisms. Arthritis Rheumatol. 2014;66(12):3337–3348.

20. Jarvela T, Paakkala T, Kannus P, Jarvinen M. The incidence of patellofemoral osteoarthritis and associated findings 7 years after anterior cruciate ligament reconstruction with bone-patellar tendon-bone autograft. Am J Sports Med. 2001;29(1):18–24.

21. Kroker A, Besler BA, Bhatla JL, et al. Longitudinal Effects of Acute Anterior Cruciate Ligament Tears on Peri-Articular Bone in Human Knees Within the First Year of Injury. J Orthop Res. 2019;37(11):2325–2336.

22. Kroker A, Bhatla JL, Emery CA, Manske SL, Boyd SK. Subchondral bone microarchitecture in ACL reconstructed knees of young women: A comparison with contralateral and uninjured control knees. Bone. 2018;111:1–8.

23. Record #57 is using an undefined reference type. If you are sure you are using the correct reference type, the template for that type will need to be set up in this output style.

24. Lohmander LS, Ostenberg A, Englund M, Roos H. High prevalence of knee osteoarthritis, pain, and functional limitations in female soccer players twelve years after anterior cruciate ligament injury. Arthritis Rheum. 2004;50(10):3145–3152.

25. Maerz T, Kurdziel M, Newton MD, et al. Subchondral and epiphyseal bone remodeling following surgical transection and noninvasive rupture of the anterior cruciate ligament as models of post-traumatic osteoarthritis. Osteoarthritis Cartilage. 2016;24(4):698–708.

26. Maerz T, Kurdziel MD, Davidson AA, et al. Biomechanical characterization of a model of noninvasive, traumatic anterior cruciate ligament injury in the rat. Ann Biomed Eng. 2015;43(10):2467–2476.

27. Maerz T, Newton M, Matthew H, Baker K. Surface roughness and thickness analysis of contrast-enhanced articular cartilage using mesh parameterization. Osteoarthritis Cartilage. 2016;24(2):290–298.

28. Maerz T, Newton MD, Fleischer M, et al. Traumatic joint injury induces acute catabolic bone turnover concurrent with articular cartilage damage in a rat model of posttraumatic osteoarthritis. J Orthop Res. 2020.

29. Maerz T, Newton MD, Kurdziel MD, et al. Articular cartilage degeneration following anterior cruciate ligament injury: a comparison of surgical transection and noninvasive rupture as preclinical models of post-traumatic osteoarthritis. Osteoarthritis Cartilage. 2016;24(11):1918–1927.

30. Mapp P, Sagar D, Ashraf S, et al. Differences in structural and pain phenotypes in the sodium monoiodoacetate and meniscal transection models of osteoarthritis. Osteoarthritis Cartilage. 2013;21(9):1336–1345.

31. McErlain D, Appleton C, Litchfield R, et al. Study of subchondral bone adaptations in a rodent surgical model of OA using in vivo micro-computed tomography. Osteoarthritis Cartilage. 2008;16(4):458–469.

32. Neuman P, Kostogiannis I, Fridén T, et al. Patellofemoral osteoarthritis 15 years after anterior cruciate ligament injury–a prospective cohort study. Osteoarthritis Cartilage. 2009;17(3):284–290.

33. Newton MD, Junginger L, Maerz T. Automated MicroCT-based bone and articular cartilage analysis using iterative shape averaging and atlas-based registration. Bone. 2020:115417.

34. Oiestad BE, Holm I, Engebretsen L, et al. The prevalence of patellofemoral osteoarthritis 12 years after anterior cruciate ligament reconstruction. Knee Surg Sports Traumatol Arthrosc. 2013;21(4):942–949.

35. Owusu-Akyaw KA, Heckelman LN, Cutcliffe HC, et al. A comparison of patellofemoral cartilage morphology and deformation in anterior cruciate ligament deficient versus uninjured knees. J Biomech. 2018;67:78–83.

36. Palmer AW, Guldberg RE, Levenston ME. Analysis of cartilage matrix fixed charge density and three-dimensional morphology via contrast-enhanced microcomputed tomography. Proc Natl Acad Sci U S A. 2006;103(51):19255–19260.

37. Potter HG, Jain SK, Ma Y, et al. Cartilage injury after acute, isolated anterior cruciate ligament tear: immediate and longitudinal effect with clinical/MRI follow-up. Am J Sports Med. 2012;40(2):276–285.

38. Ramme A, Lendhey M, Raya J, Kirsch T, Kennedy O. A novel rat model for subchondral microdamage in acute knee injury: a potential mechanism in post-traumatic osteoarthritis. Osteoarthritis Cartilage. 2016;24(10):1776–1785.

39. Stoop R, Buma P, van der Kraan PM, et al. Type II collagen degradation in articular cartilage fibrillation after anterior cruciate ligament transection in rats. Osteoarthritis Cartilage. 2001;9(4):308–315.

40. Tsai PH, Lee HS, Siow TY, et al. Abnormal perfusion in patellofemoral subchondral bone marrow in the rat anterior cruciate ligament transection model of post-traumatic osteoarthritis: a dynamic contrast-enhanced magnetic resonance imaging study. Osteoarthritis Cartilage. 2016;24(1):129–133.

41. van Meer BL, Waarsing JH, van Eijsden WA, et al. Bone mineral density changes in the knee following anterior cruciate ligament rupture. Osteoarthritis Cartilage. 2014;22(1):154–161.

