## Supplementary material for "Early Patellofemoral Cartilage and Bone Degeneration in a Rat Model of Noninvasive Anterior Cruciate Ligament Rupture": IACUC Approval

Research Institute Beaumont | HEALTH SYSTEM

TO: Kevin Baker, Ph.D.
Orthopaedic Research

FROM: Brian Marples, Ph.D.
Chairman, Animal Care Committee

DATE: November 18, 2013

SUBJECT: **Status of Application AL-13-14 (Full Approval)**
"Systemic and Local Responses to Anterior Cruciate Ligament Rupture"

Thank you for submitting your revised protocol clarifying the minor concerns raised by the Animal Care Committee (ACC) during the review process of your protocol. I am pleased to inform you that your protocol was granted **full approval**.

This approval is in effect for one year. You are required to submit a progress report annually or upon project completion, whichever comes first. If continuation of a project is desired beyond the initial approval period, you may request a one-year continuation when completing the "Annual Review" form. Approval for continuation of a project may be granted twice. The maximum approval period for any one project is three years.

In addition, **schedules cannot be confirmed until** all associates listed on the protocol who are identified as working on this study fulfill the mandatory William Beaumont Hospital Research Services Training Program.

1. Review "Research Services Training Manual" and acknowledge understanding by signature; the signature form is enclosed and an electronic copy of the manual may be accessed on "Inside Beaumont" at http://employee.beaumont.edu/portal/pls/portal/docs/1122883.PDF
2. Complete both CITI (Collaborative Institutional Training Initiative) web-based courses:
   a. "Working with the IACUC for Investigators, Staff and Students"
   b. "Working with Rats in Research Setting"

Federal policy requires prior Committee review and approval before any significant changes to an approved protocol are initiated. Discrepancies between protocol statements and actual animal use may result in delay or withdrawal of approval. Examples of significant changes are described in policy #404 "Significant Changes to Approved Animal Research Protocols". This and other ACC polices may be accessed on Inside Beaumont online at http://employee.beaumonthospitals.com/portal/pls/portal/docs/1042466.PDF or by contacting the ACC office.

The *Guide for the Care and Use of Laboratory Animals, 8$^{th}$ Edition* is the primary reference on standards of animal care used at William Beaumont Hospital. Our institutional assurance statement, #A3408-1, has been approved through September 30, 2017 by the Office for Laboratory Animal Welfare (OLAW). In addition, we have been awarded Full Accreditation by the Association for Assessment and Accreditation of Laboratory Animal Care (AAALAC International).

cc: Kyle Anderson, M.D.
Joel Hein, M.D.
Michele McGonagle, B.I.S., L.V.T., L.A.T.
Katie Golden-Prehm
Brett Habermas

BM/em
