## Supplementary material for "Early Patellofemoral Cartilage and Bone Degeneration in a Rat Model of Noninvasive Anterior Cruciate Ligament Rupture": ARRIVE Checklist

### The ARRIVE Guidelines Checklist

#### Animal Research: Reporting In Vivo Experiments

Carol Kilkenny[1], William J Browne[2], Innes C Cuthill[3], Michael Emerson[4] and Douglas G Altman[5]

*[1]The National Centre for the Replacement, Refinement and Reduction of Animals in Research, London, UK, [2]School of Veterinary Science, University of Bristol, Bristol, UK, [3]School of Biological Sciences, University of Bristol, Bristol, UK, [4]National Heart and Lung Institute, Imperial College London, UK, [5]Centre for Statistics in Medicine, University of Oxford, Oxford, UK.*

| | ITEM | RECOMMENDATION | Section/ Paragraph |
|---|---|---|---|
| Title | 1 | Provide as accurate and concise a description of the content of the article as possible. | Title |
| Abstract | 2 | Provide an accurate summary of the background, research objectives, including details of the species or strain of animal used, key methods, principal findings and conclusions of the study. | Abstract |
| **INTRODUCTION** | | | |
| Background | 3 | a. Include sufficient scientific background (including relevant references to previous work) to understand the motivation and context for the study, and explain the experimental approach and rationale.<br>b. Explain how and why the animal species and model being used can address the scientific objectives and, where appropriate, the study's relevance to human biology. | a. Intro: P1-2<br>b. Intro: P3 |
| Objectives | 4 | Clearly describe the primary and any secondary objectives of the study, or specific hypotheses being tested. | Intro: P3 |
| **METHODS** | | | |
| Ethical statement | 5 | Indicate the nature of the ethical review permissions, relevant licences (e.g. Animal [Scientific Procedures] Act 1986), and national or institutional guidelines for the care and use of animals, that cover the research. | Mthds: P1 |
| Study design | 6 | For each experiment, give brief details of the study design including:<br>a. The number of experimental and control groups.<br>b. Any steps taken to minimise the effects of subjective bias when allocating animals to treatment (e.g. randomisation procedure) and when assessing results (e.g. if done, describe who was blinded and when).<br>c. The experimental unit (e.g. a single animal, group or cage of animals).<br>A time-line diagram or flow chart can be useful to illustrate how complex study designs were carried out. | a. Mthds: P1<br>b. Mthds: P1, P4<br>c. Mthds: P1 |
| Experimental procedures | 7 | For each experiment and each experimental group, including controls, provide precise details of all procedures carried out. For example:<br>a. How (e.g. drug formulation and dose, site and route of administration, anaesthesia and analgesia used [including monitoring], surgical procedure, method of euthanasia). Provide details of any specialist equipment used, including supplier(s).<br>b. When (e.g. time of day).<br>c. Where (e.g. home cage, laboratory, water maze).<br>d. Why (e.g. rationale for choice of specific anaesthetic, route of administration, drug dose used). | a. Mthds: P1<br>b. Mthds: P1<br>c. Mthds: P1<br>d. Mthds: P1 |
| Experimental animals | 8 | a. Provide details of the animals used, including species, strain, sex, developmental stage (e.g. mean or median age plus age range) and weight (e.g. mean or median weight plus weight range).<br>b. Provide further relevant information such as the source of animals, international strain nomenclature, genetic modification status (e.g. knock-out or transgenic), genotype, health/immune status, drug or test naïve, previous procedures, etc. | a. Mthds: P1<br>b. Mthds: P1 |

| | | | |
|---|---|---|---|
| Housing and husbandry | 9 | Provide details of:<br>a. Housing (type of facility e.g. specific pathogen free [SPF]; type of cage or housing; bedding material; number of cage companions; tank shape and material etc. for fish).<br>b. Husbandry conditions (e.g. breeding programme, light/dark cycle, temperature, quality of water etc for fish, type of food, access to food and water, environmental enrichment).<br>c. Welfare-related assessments and interventions that were carried out prior to, during, or after the experiment. | a. Mthds: P1<br>b. Mthds: P1<br>c. Results: P1 |
| Sample size | 10 | a. Specify the total number of animals used in each experiment, and the number of animals in each experimental group.<br>b. Explain how the number of animals was arrived at. Provide details of any sample size calculation used.<br>c. Indicate the number of independent replications of each experiment, if relevant. | a. Mthds: P1<br>b. Mthds: P1<br>c. N/A |
| Allocating animals to experimental groups | 11 | a. Give full details of how animals were allocated to experimental groups, including randomisation or matching if done.<br>b. Describe the order in which the animals in the different experimental groups were treated and assessed. | a. Mthds: P1<br>b. Mthds: P1 |
| Experimental outcomes | 12 | Clearly define the primary and secondary experimental outcomes assessed (e.g. cell death, molecular markers, behavioural changes). | Mthds: P2-4 |
| Statistical methods | 13 | a. Provide details of the statistical methods used for each analysis.<br>b. Specify the unit of analysis for each dataset (e.g. single animal, group of animals, single neuron).<br>c. Describe any methods used to assess whether the data met the assumptions of the statistical approach. | a. Mthds: P5<br>b. Mthds: P5<br>c. Mthds: P5 |
| RESULTS | | | |
| Baseline data | 14 | For each experimental group, report relevant characteristics and health status of animals (e.g. weight, microbiological status, and drug or test naïve) prior to treatment or testing. (This information can often be tabulated). | Methods: P1 |
| Numbers analysed | 15 | a. Report the number of animals in each group included in each analysis. Report absolute numbers (e.g. 10/20, not 50%[2]).<br>b. If any animals or data were not included in the analysis, explain why. | a. Results: P1<br>b. N/A |
| Outcomes and estimation | 16 | Report the results for each analysis carried out, with a measure of precision (e.g. standard error or confidence interval). | Results: P2-4 |
| Adverse events | 17 | a. Give details of all important adverse events in each experimental group.<br>b. Describe any modifications to the experimental protocols made to reduce adverse events. | a. N/A<br>b. N/A |
| DISCUSSION | | | |
| Interpretation/ scientific implications | 18 | a. Interpret the results, taking into account the study objectives and hypotheses, current theory and other relevant studies in the literature.<br>b. Comment on the study limitations including any potential sources of bias, any limitations of the animal model, and the imprecision associated with the results[2].<br>c. Describe any implications of your experimental methods or findings for the replacement, refinement or reduction (the 3Rs) of the use of animals in research. | a. Disc: P1-4<br>b. Disc: P5<br>c. Disc: P5 |
| Generalisability/ translation | 19 | Comment on whether, and how, the findings of this study are likely to translate to other species or systems, including any relevance to human biology. | Disc: P3 |
| Funding | 20 | List all funding sources (including grant number) and the role of the funder(s) in the study. | Acknowledgm ents |


References:

1. Kilkenny C, Browne WJ, Cuthill IC, Emerson M, Altman DG (2010) Improving Bioscience Research Reporting: The ARRIVE Guidelines for Reporting Animal Research. *PLoS Biol* 8(6): e1000412. doi:10.1371/journal.pbio.1000412
2. Schulz KF, Altman DG, Moher D, the CONSORT Group (2010) CONSORT 2010 Statement: updated guidelines for reporting parallel group randomised trials. *BMJ* 340:c332.
